# ChemBERTaPolyPharm: Modeling polypharmacy side effects with ChemBERTa and PubMed Encoders

**DOI:** 10.1101/2025.05.20.655109

**Authors:** Anastasiya A. Gromova, Anthony S. Maida

## Abstract

Polypharmacy side effects occur when drug combinations trigger unexpected interactions, altering therapeutic outcomes. During which the activity of one drug may change favorably or unfavorably if taken with the other drug. Drug interactions are rare and are only observed in clinical studies. Thus, the discovery and detection of polypharmacy side effects remains a challenge. Our approach achieves an impressive average F1 score of 0.93, demonstrating its efficacy in capturing drug interaction patterns.

**Submission of papers to NeurIPS 2025:** Please read the instructions below carefully and follow them faithfully.

## 1.1 Introduction

The term polypharmacy refers to the simultaneous use of multiple medications to treat patients with complex health conditions. Typically, a drug combination consists of multiple drugs, where each individual drug has been validated as a single effective medication treatment within the patient population. The polypharmacy side effect problem arises when two or more drugs taken in combination cause a negative reaction or adverse side effects, even when the use of the drugs individually causes no harm. Drug-drug interactions (DDIs) are a major cause of these adverse reactions, and predicting potential DDIs is becoming increasingly important due to the growing number of possible prescribed drug combinations. Adverse side effects often arise due to drug-drug interactions (DDIs), where two or more medications, taken simultaneously, interact with each other, potentially impeding, augmenting, or diminishing the intended effects of a drug, or in more serious scenarios, triggering undesirable side effects. An adverse drug reaction (ADR) is a harmful or unpleasant response resulting from the use of a medicinal product. ADRs can significantly affect patient safety, treatment effectiveness, and overall health outcomes. Importantly, ADRs resulting from DDIs can range from symptoms of mild discomfort to life-threatening consequences. Polypharmacy side effects are unique in that they are not directly associated with either of the individual drugs within the pair. Detecting all possible DDIs during clinical trials in a laboratory setting is impractical and costly. Typically, lab clinical trials focus on studying the individual side effects of a drug rather than its interactions in combination with other medications. Various computational methods have been proposed to address this challenge. However, many rely on external biomedical knowledge, limiting their applicability for the prediction of potential DDIs with new drugs. The prevalence of polypharmacy among chronically ill adults is high due to the prevalence of multimorbidity, where individuals are affected by multiple health conditions. According to the U.S. Centers for Disease Control and Prevention (CDC), over 10% of individuals use five or more drugs concurrently, with 20% of older adults taking at least ten drugs [1]. With an aging population experiencing prevalence of chronic conditions in the United States (U.S.), a significant portion of these patients are on multiple medications. High-risk medications can increase the risk for DDIs [2]. The World Health Organization (WHO) emphasizes the gravity of medication safety in polypharmacy. Their report highlights that much of the harm caused by ADRs is preventable. Surprisingly, adverse events now rank as the fourteenth leading cause of morbidity and mortality worldwide, placing patient harm on par with diseases like tuberculosis and malaria [3]. As the number of approved drugs continues to rise, the likelihood of interactions increases [4]. One approach to mitigate these risks is to conduct small pre-clinical in vitro safety trials to detect DDIs. These trials involve testing drug combinations in a controlled laboratory environment to observe any adverse interactions before proceeding to clinical trials. However, these trials are limited in scale, require a long duration, and are costly [5]. The task of manually identifying polypharmacy side effects is complex, and with the number of potential drug combinations, it becomes practically impossible to test directly. Furthermore, the manual detection of polypharmacy side effects is challenging [6], [7]. In 2002, it was estimated that polypharmacy incurred an annual cost of at least 50 billion for U.S. healthcare plans, representing a significant share of pharmaceutical expenditure [8]. According to Ernst and Grizzle [9], the annual cost of polypharmacy treatment in the U.S. alone exceeds 177 billion. From 1999-2000 to 2017-2018, the overall percentage of adults with polypharmacy consistently increased. Specifically, with the upward rising trend from 8.2% to 17.1% highlights the challenge of manually identifying polypharmacy side effects, as they are rare and typically not observed in small clinical trials [8]. Consequently, many ADRs remain undiscovered. Given that the manual clinical methods to perform systematic combinatorial screening are costly and labor-intensive [5]-[9], there is a clear need for more efficient alternatives. As a result, predicting and managing polypharmacy becomes an urgent imperative in clinical practice. The rising number of approved drugs highlights the need for robust polypharmacy and DDI prediction in patient treatment plans and drug design processes, emphasizing the importance of both patient safety and pharmaceutical development. To address this need, there has been a significant shift towards computational methods, particularly machine and deep learning, to predict DDIs and their associated adverse ADRs. Despite advancements in deep learning, predicting DDIs remains a complex and challenging task. Traditional methods for identifying drug-drug interactions (DDIs) focus on text-mining, a key task within Natural Language Processing (NLP). With the growth of biomedical literature, a significant amount of drug-related knowledge is hidden in unstructured texts such as published articles, scientific journals, books, and technical reports. Text-mining techniques, particularly relation extraction, are designed to detect DDIs from the textual information [10], [11], [12], [13], [14], [15], [16], [17], [18]. This process transforms the DDI problem into a relationship extraction task, where the goal is to identify and classify interactions between drugs based on their mentions in the literature. Relation extraction aims to identify specific relationships between entity pairs (such as drug pairs) mentioned in documents. Researchers have compiled and annotated these interactions to create DDI databases. However, these methods have limitations, as they often fail to identify unannotated DDIs and flag potential interactions before combinational treatments. Recently, computational approaches have been increasingly adopted to analyze DDIs from both machine learning and molecular mechanisms perspectives. These methods offer a promising avenue for identifying previously unannotated potential DDIs. The use of neural networks and graph convolutional neural networks has shown promise in modeling and predicting DDIs by learning from historical data on drug co-occurrences and their effects. Through the neural-network-based approaches and systems, researchers construct comprehensive interaction networks incorporating drugs, targets, and pathways, effectively capturing topological relationships among drugs. Integrating graph-based machine learning approaches helps uncover hidden interactions and identify novel targets for mitigating DDIs. Computational machine learning methods offer a robust and innovative approach to addressing the intricate challenge of drug-drug interactions (DDIs). By integrating diverse datasets that encompass molecular structures, biological pathways, protein interactions, and target information from both biomedical and clinical domains, these advanced techniques can unravel the complex mechanisms underlying DDIs. This analysis not only enhances our understanding of how different drugs interact at a chemical structure level but also sheds light on the broader implications for patient safety and therapeutic efficacy. Consequently, machine learning models are becoming indispensable tools in the field of pharmacology, aiding in the prediction, prevention, and management of adverse drug interactions. Given the intricacies of polypharmacy, these insights are crucial for addressing its two primary challenges (two-fold problem):

1. Prediction of Drug-Drug Interactions: Determining potential interactions between two drugs.
2. Side Effect Manifestation: Identifying the specific types of side effects that may occur due to these interactions.

The key takeaways from our comprehensive experiments are as follows: This work explores how to incorporate domain knowledge available in form of chemical molecular structures of drugs and pubmed corpus embeddings combined with the deep learning techniques to predict drug-drug interactions (DDIs), which are critical for patient safety and effective treatment plan. Our main research questions are as follows:

- Given the high cost and time associated with clinical trials and the difficulty in manually identifying polypharmacy side effects, what alternative sources of information as drug characteristics can be utilized to address the polypharmacy problem for new pairs of drugs where clinical studies may not yet be available?
- How can domain knowledge, specifically in the form of molecular structures of drugs and pubmed mono side effects features be effectively incorporated to enhance the prediction of DDI?
- How can we develop a robust framework to predict DDIs for new pairs of drugs that were not included in the training set?

By incorporating data on chemical molecular structures, biological pathways, protein, and targets from biomedical and clinical domains, these techniques provide valuable insights into mechanisms and implications of DDIs. This comprehensive approach not only enhances our understanding of drug interactions but also contributes to the development of safer and more effective therapeutic strategies. In this paper, we address the complex issue of polypharmacy, particularly under the constraint that clinical trials are both costly and time-consuming. Identifying polypharmacy side effects manually is challenging, especially for new drug pairs where clinical studies may not be readily available. To tackle this problem, we explore alternative sources of information that can aid in predicting drug-drug interactions (DDIs) for these new pairs. Our approach leverages domain knowledge encapsulated in the chemical molecular structure of drugs to enhance DDI prediction. We introduce a novel method, ChemBERTa-DDI, which integrates drug embeddings derived from the rich chemical structure of drugs (ChemBERTa) with a deep learning neural network (DNN) model serving as the predictor. Specifically, we augment the drug data with external chemical structure information in the form of the drug’s canonical SMILES (Simplified Molecular Input Line Entry System) obtained from PubChem. The DNN predictor operates by concatenating the feature vectors of any two drugs to form a concatenated feature vector representing the corresponding drug pair. This composite vector is then used to train the DNN to predict potential DDIs. Our method not only utilizes the inherent chemical properties of the drugs but also incorporates advanced machine learning techniques to improve prediction accuracy. To validate our approach, we compare the performance of various machine learning classifiers, including XGBoost, in predicting DDIs using the drug pair features generated by our method. Experiments conducted on the TWOSIDES dataset demonstrate that our strategy significantly outperforms other strong baseline architectures, achieving an impressive 0.93 F1 score. Moreover, our model exhibits the capability to predict potential DDIs for new compounds that were not included in the training set. This innovation extends the applicability of our approach to real-world scenarios where new drugs are continually being developed and introduced. By comparing our model with established methods, we provide a comprehensive evaluation of its performance in predicting DDIs, highlighting its potential to enhance drug safety and efficacy in clinical practice.

## 2. Literature Review

Traditional approaches to predicting drug-drug interactions (DDIs) fall into two main categories: text mining-based and machine-learning based approaches. Text mining-based methods [10], [11], [12], [13], [14], [15], [16], [17], [18] extract annotated DDIs from electronic medical records and scientific literature. These methods are valuable for constructing DDI-related databases. However, they have significant limitations, the inability to detect unannotated DDIs and to provide alerts for potential DDIs before combinational treatments are administered in real-world scenarios. Machine learning based approaches build prediction models based on the known DDIs in databases and predict novel ones. To create a landscape of DDI methods, we can organize the DDI prediction methods into two categories: Literature-based and Machine Learning/ Deep Learning Approaches:

### 2.1 Literature-Based Learning Approaches

#### 2.1.1 NLP-Based Literature Extraction

Literature-based extraction methods utilize natural language processing (NLP) techniques to extract drug-drug interactions (DDIs) from biomedical literature [10], [12], [13], [14], [15], [16], [17]. DDI extraction is generally regarded as a Relation Extraction task (RE), aiming to identify specific relationships between entity pairs within documents that mention these pairs. The DDIExtraction challenges organized in 2011 [12] and 2013 [13], [10] provided annotated corpora for training and testing purposes. Notably, the 2013 DDIExtraction task required participants to identify DDIs and classify them into specific types, effectively modeling the task as a multi-class classification problem. These challenges have significantly contributed to advancing the field by providing standardized datasets and evaluation metrics.

#### Conventional Classifier Approaches

These methods utilize hand-crafted features to extract drug-drug interactions (DDIs) from biomedical literature. For example, Tari et al. [11] employ hand-made features and reasoning based on drug metabolism properties to identify DDIs in biomedical texts. However, this approach leverages domain-specific knowledge, which can be labor-intensive and may not generalize well to new data without extensive feature engineering.

### 2.2 Machine Learning (Semi-Supervised) Learning Approaches

#### 2.3 Classification-based approaches

The traditional classification-based method transforms the drug-drug interaction prediction task into a binary classification problem. It uses labeled data of interacting and non-interacting drug pairs to train models that predict interactions for each side effect type. These approaches employ various machine learning models to classify and predict whether two drugs interact. Conventional classifiers such as Support Vector Machines (SVM) and decision trees are commonly used for DDI prediction. For instance, Ferdousi et al. [18] utilize functional similarities between drugs for computational prediction of DDIs. Similarly, Qian et al. [19] leverage genetic interactions to predict adverse DDIs.

#### 2.4 NN-based Approaches

These approaches use neural networks (NN) to construct advanced and expressive features for predicting DDIs ([20], [21], [22], [23], [24], [25], [26], [27], [28], [29]). Examples include methods like Decagon [20], NNPS [21], DPSP [22], DPDDI [24], DeepDDI [25], DSN-DDI [26], De-SIDEDDI [27], and purely SMILES-based model [28]. Zitnik et al. [20] model polypharmacy side effects using graph convolutional networks Decagon, while Masumshah et al. [21] employ neural networks for polypharmacy side effects prediction using PCA representations of drug-mono side effects and drug-protein representation. Later, Masumshah et al. [22] introduced DPSP, which combines Jaccard similarity with deep neural networks. Bumgardner et al. [28] propose a novel method for predicting DDIs based on the vital chemical substructure of drugs extracted from their SMILES strings. They construct a graph that connects drugs based on their common functional chemical substructures and apply graph neural network (GNN) methods to generate drug embeddings. These embeddings are then used to predict DDIs.

#### 2.5 Similarity-based Approaches

These approaches are founded on the principle that drugs with similar chemical structures or properties are likely to interact with the same drugs, as highlighted by Vilar et al [30]. To predict drug-drug interactions (DDIs), these methods utilize similarity measures between drugs, often incorporating domain-specific, hand-crafted features to enhance prediction accuracy. However, many of these approaches are limited by the use of small datasets and a narrower focus into account limited datasets and fewer drug-centric interactions.

In the realm of drug-drug interactions (DDIs), most machine learning-based approaches follow a typical workflow that usually contains two key components.

1. **Feature Extraction:** This feature extraction component is essential for transforming raw data into meaningful features that can be utilized for accurate predictions. The feature extraction process converts drug properties into feature vectors, which may include chemical structures, targets, Anatomical Therapeutic Chemical (ATC) codes, side effects, and clinical observations.
2. **Supervised Prediction:** Utilizing the features extracted, this component trains machine learning models to predict potential DDIs. The supervised prediction component employs various classification algorithms such as KNN, SVM, logistic regression, decision trees, and naïve Bayes, as well as network propagation methods like reasoning over drug-drug network structures, label propagation, random walks, probabilistic soft logic, and matrix factorization. The predictor is trained using both feature vectors/similarity matrices and annotated DDI labels to identify potential DDIs. While most methods use a single predictor, some integrate multiple predictors for enhanced accuracy.

By integrating these components, machine learning-based approaches can effectively leverage diverse data types and sophisticated algorithms to enhance the accuracy and reliability of DDI predictions.

## 3 Methodology

Many studies address the polypharmacy problem as two-fold drug side effect prediction. Recently, the new popular feature extraction method in the field of drug development and discovery is the Graph Convolution Network (GCN). GCNs perform convolution on graphs, aggregating information from each node’s neighborhood to create dense vector embeddings. These low dimensional representation embeddings capture the structural relationships between nodes (drugs) in the network without requiring manual feature engineering. These dense vector embeddings can be used as features in machine learning models for downstream tasks, such as link prediction. Link prediction is a task in graph theory and network analysis where the goal is to predict the existence of a link between two nodes in a network. In the context of drug side effect prediction, link prediction can help identify potential interactions between drugs that have not been previously documented, thereby aiding in the discovery of new side effects or therapeutic uses. The core paper, Zitnick et al. [20], introduced Decagon which is a model uses GCN for multirelational link prediction in heterogeneous graphs. It performs semi-supervised learning on graphs. It is a multi-relational model, meaning it can handle multiple types of relationships in the data. This is achieved by constructing a multimodal graph with three edge types to encode protein-protein interactions, drug-protein targets, and drug-drug interactions. This information is obtained by using a catalog of 964 different polypharmacy side effects. Decagon is composed of two components: an encoder, which is a GCN that operates on the graph and produces embeddings for nodes, and a decoder, which is a tensor factorization model which uses embeddings to model polypharmacy side effects. Decagon performs multirelational link prediction which means that it tries to identify not yet observed true links. For link prediction , Decagon uses a GCN model, which involves predicting associations between pairs of drugs and potential side effects ‘drug side effect prediction. Decagon models particularly well, polypharmacy side effects that have a strong molecular basis, leveraging dependencies between side effects and reusing the information learned about the molecular basis of one side effect to better understand the molecular basis of another side effect. The study by Masumshah et al. [21], innovatively transforms the polypharmacy DDI prediction problem into a simple binary matrix problem. The neural network-based method for polypharmacy side effects prediction (NNPS), uses a neural network, combined with an approach to feature representation to predict DDIs. Initially, feature extraction is performed to create feature vectors for drugs based on their mono side effects and drug-protein interaction information. To enhance computational efficiency and reduce dimensionality, dimensionality reduction is applied using Principal Component Analysis (PCA) to drug mono side effects and drug protein information, resulting in two reduced feature matrices to retain 95% of the original variance. For a given drug, feature vectors are created that include 503 features for mono side effects and 22 features for drug-protein interactions. These matrices are concatenated into a single feature vector of length 523 for each drug. For a given drug pair, (drug_i,drug_j), the feature aggregation is performed where corresponding feature vectors are concatenated. The model constructs a separate binary matrix for each of the 964 side effects, with ground truth data indicating whether each drug pair causes that side effect. The neural network is trained independently for each side effect, resulting in 964 specialized models. Each model outputs the probability of a side effect occurring, based on a predetermined threshold. This approach not only enhances the accuracy of DDI predictions but also provides a scalable solution for managing the complexity of polypharmacy side effects. In the latest study by Masumshah et al. [22], titled “DPSP: A Multimodal Deep Learning Framework for Polypharmacy Side Effect Prediction”, advance their previous work by introducing a more enhanced framework for predicting polypharmacy side effects. As stated by the authors, NNPS [21] “NNPS only uses a small number of features and may not capture the full complexity of DDI”. In contrast, the DPSP framework uses feature extraction methods and the Jaccard similarity to determine similarities between drugs, generating novel feature vectors. The study by Yue-Hua Feng et al. [24] proposes a method called DPDDI to predict DDIs. This method works by extracting the low-dimensional feature representation structure features of drugs in a graph embedding space, capturing topological relationship to their neighborhood drugs of each drug from DDI network using GCN and feeding these features into a Deep Neural Network (DNN) model for prediction. The DPDDI method consists of three phases: Firstly, the GCN model is used to extract the low-dimensional embedding features of drugs from DDI network. Secondly, aggregate the extracted structure network latent features of drugs for the drug pairs. The authors experimented with various aggregation operators and found that concatenation yields best results. Finally, the concatenated drug pair feature vectors are fed into a DNN to predict potential DDIs. The strength of the DPDDI method lies in its ability to extract network structure features, effectively capturing topological relationships between drugs. DPDDI is effective in predicting potential interactions between drugs present in the DDI network. However, it fails when the DDI network does not include certain drugs, such as newly invented drugs without prior information. From the literature-based method perspective, the paper by Mondal et al [17] creates BERTChem-DDI, a novel method for predicting DDIs by leveraging both textual data and chemical structure information. The method combines drug embeddings derived from the chemical structures of drugs Simplified Molecular Input Line Entry System (SMILES), with BioBERT embeddings, which are pre-trained on biomedical text. The approach integrates domain-specific knowledge into the RE task, enhancing the prediction accuracy of DDIs from textual data. The combination of chemical and textual data represents a significant advancement in the field of biomedical/pharmacological natural language processing.

**Table 1.** Pre-training Datasets.

| Datasets |  |
| --- | --- |
| Name | Description |
| PubMed | Input Medical Articles Corpus |
| SMILES | Input Molecular Structure Corpus |
| Protein | Input Protein Information Corpus |

**Table 2.** Databases details.

| Database | Details |
| --- | --- |
| **TWOSIDES** | No. of side effects = 1,317<br>No. of drugs = 645<br>No. of drug pairs = 63,473 |
| **SIDER, OFF-SIDES** | No. of SIDER drugs = 1,556<br>No. of SIDER side effects = 5868<br>No. of OFFSIDES drugs = 1,332<br>No. of OFFSIDES side effects = 10,097 |
| **STITCH** | No. of compounds = 519,022<br>No. of protein targets = 8,934<br>No. of interactions = 8,083,600 |
| **PubChem** | No. of SMILES = 645 |

### 3.1 Model Training

The pretraining was performed using two Encoder models ChemBERTa (MLM) 77M parameters and PubMed base (MLM). Text and word tokenization was performed using AutoTokenizer from the HuggingFace ‘Autotokenizer’. The maximum sequence length is 1024. Training was performed using the ‘Standard NC80 H100’ compute node. In our previous work ChemBERTaDDI [23], Gromova outlined the approach using ChemBERTa encoder. Building on top of the ChemBERTa and PubMed Encoders, we are going to focus on vectorization of the molecular input (SMILES) and the PubMed corpus for the mono-side effects as the inputs.

#### Feature Extraction

The AutoTokenizer tokenizer converts SMILES strings into token IDs, which are then fed into the *ChemBERTa-77M-MLM* transformer model developed by Chithranda et al. [31]. The advancements in self-supervised pretraining for molecular property prediction *ChemBERTaDDI* [22], which employs *ChemBERTa-77M-MLM* embeddings derived from transformer attention mechanisms. The *ChemBERTa* model processes tokens through multiple encoder layers to generate contextualized molecular embeddings. Serving as a robust feature extractor, *ChemBERTa-77M-MLM* transforms SMILES inputs into generalizable embeddings that effectively capture the chemical structure and properties of each drug based on its molecular composition. In addition, PubMed corpus is leveraged as base model from the mono side effects description of side efffect PubMed is kept as a frozen encoder that provides fixed emebeddings of size 384.

#### Feature Aggregation

The PCA-reduced clinical mono side effect features are concatenated with *ChemBERTa* embeddings to form a unified 979-dimensional feature vector for each drug. Subsequently, for each drug pair, their respective individual feature vectors are aggregated using an element-wise summation operation. The summation gate effectively combines the attributes of both drugs, resulting in a comprehensive feature vector that encapsulates the joint clinical and chemical characteristics necessary for predicting DDIs. For each side effect the corresponding PubMedBERT feature is computed and aggregated across drugs using the binary incidence matrix defined by clinical drug–side–effect associations. To reduce dimensionality principal component analysis (PCA) is applied to the aggregated embeddings, yielding a compact representation where *n_components* 384 components that preserves essential variational information.

### 3.2 Experiments and Results

#### 3.2.1 Polypharmacy Prediction (Drug-to-drug interaction (DDI) Task)

Downstream results analysis comparing the models trained on identical datasets. Our experimental results demonstrate that ChemBERTaDDI achieves an average F1 score of 0.94, while our proposed ChemBERTaPolyPharm—integrating both PubMedBERT and ChemBERTa encoders—attains an average F1 score of 0.93. This clearly demonstrates that ChemBERTaPolyPharm contains both embeddings PubMed and ChemBERTa encoders. For the architectural efficiency and consistency we keep using the same feedforward NN as performed in previous work ChemBERTaDDI [23] across predicting 964 polypharmacy tasks.

## 4 Feed-Forward Neural Network Architecture

The feedforward NN architecture is comprised of following three-layers:

- First Hidden Layer: A dense layer with 250 neurons, initialized with the Glorot. This is followed by batch normalization, a ReLU activation, and a dropout layer with a rate of 0.3.
- Second Hidden Layer: A dense layer containing 300 neurons with Glorot initialization, batch normalization, ReLU activation, and a dropout of 0.3.
- Third Hidden Layer: A dense layer with 200 neurons, also using Glorot initialization, followed by batch normalization, ReLU activation, and a dropout rate of 0.3.

## 5 Results

In this section, we present a comprehensive analysis of the experimental results and findings obtained during our study. The comparison is performed for the ChemBERTaPolypharm with five well-established methods: Decagon[1], NNPS [2], DeepWalk [11], DEDICOM [12], and RESCAL [13]. The performance metrics for each method areas follows: Decagon achieved an ROC of 0.874 and an AUPR of 0.825. DeepWalk attained an ROC of 0.761 and an AUPR of 0.737. DEDICOM obtained an ROC of 0.705 and an AUPR of 0.637.RESCAL reached an ROC of 0.693 and an AUPR of 0.613. NNPS achieved an ROC of 0.966 and an AUPR of 0.953. Our experiments demonstrate that the ChemBERTaPolypharm and ChemBERTaDDI significantly outperforms these baseline methods, as shown in Table III, achieving an F1-score of 0.93 and 0.94 respectively, with AUPR of 0.95, and an AUROC of 0.97. In addition, Table IV presents the results of a case study evaluating the performance of ChemBERTaPolypharm and ChemBERTaDDI against Decagon in predicting dangerous polypharmacy side effects. The evaluation is measured by the Area Under the Precision-Recall Curve (AUPR) and includes supporting literature evidence. Across all examined conditions, ChemBERTaPolypharm and ChemBERTaDDI consistently demonstrate superior predictive accuracy. Our findings show that by incorporating PubMed corpus with mono side effects descriptions and ChemBERTa we have achieved a significantly better performance on the downstream task.

**Figure 1.**
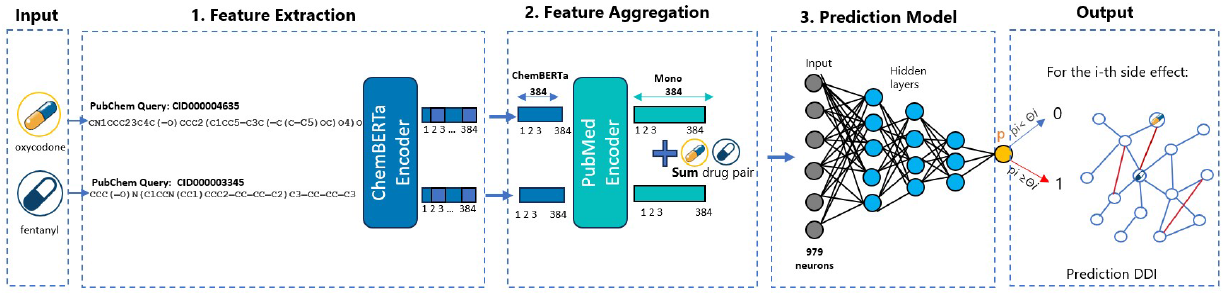
ChemBERTaPolyPharm: Modeling polypharmacy side effects with ChemBERTa and PubMed Encoder.

**Figure 2.**
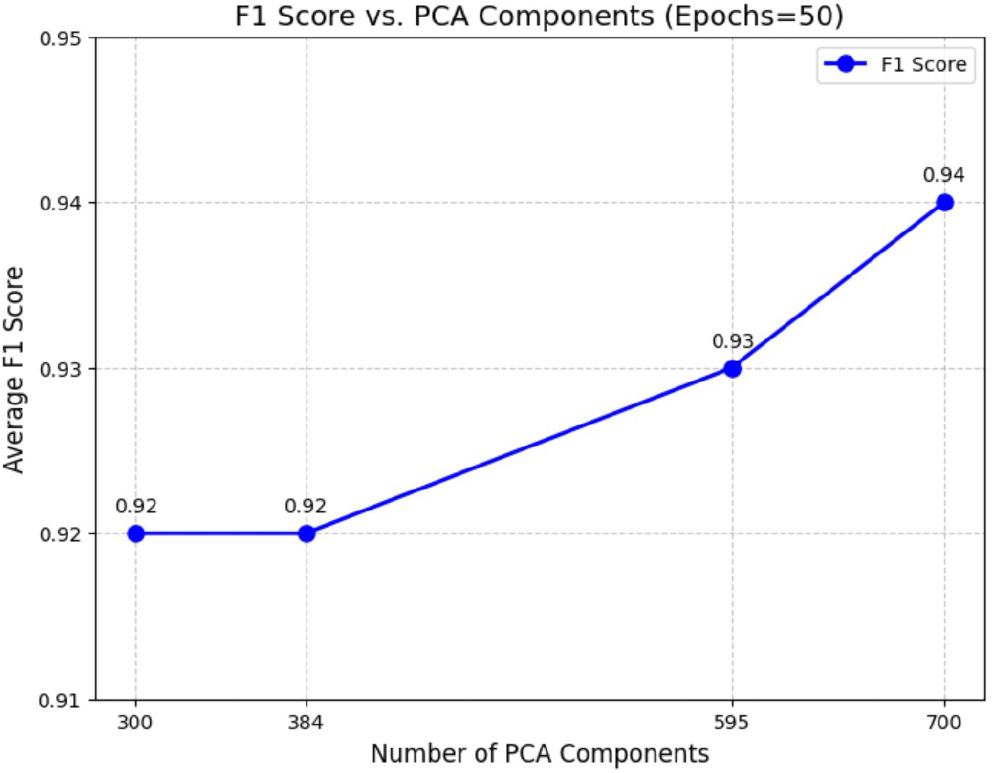
ChemBERTaPolyPharm:Average F1 score.

**Table 3.** Model Downstream Evaluation Task Results.

| Method | ROC | AUPR | Recall | Precision | F1 Score |
| --- | --- | --- | --- | --- | --- |
| NNPS | 0.966 | 0.530 | 0.976 | 0.901 | 0.936 |
| Decagon | 0.874 | 0.825 | 0.950 | 0.771 | 0.850 |
| ChemBERTaDDI | 0.965 | 0.954 | 0.968 | 0.909 | 0.937 |
| ChemBERTaPolyPharm | 0.958 | 0.947 | 0.959 | 0.893 | 0.930 |

**Table 4.** Case Study: Results of Dangerous Polypharmacy Side Effects in ChemBERTaDDI and Decagon on AUPR.

| Polypharmacy | AUPR (ChemBERTaDDI) | AUPR (Decagon) | Evidence |
| --- | --- | --- | --- |
| Sarcoma | 0.978 | 0.789 | Serban et al. [32] |
| Carcinoma of the Cervix | 0.952 | 0.810 | Arbyn et al. [33] |
| Malignant Hypertension | 1.00 | 0.858 | Januszewicz et al. [34] |
| Oophorectomy | 0.974 | 0.911 | Evans et al. [35] |
| Leukopenia | 0.960 | 0.695 | Flanagan et al. [36] |

## 6 Conclusion

This paper developed a new computational ChemBERTaPolypharm along with previously developed ChemBERTaDDI framework for drug-drug interaction prediction that effectively integrates high-dimensional clinical data with the PubMed corpus with transformer-based chemical molecular embeddings. The ChemBERTaPolyPharm framework leverages a SMILES-based ChemBERTa and PubMed encoders to extract a 384-dimensional embedding, which is then concatenated with a 384-dimensional mono side-effect feature vector before being passed into a feed-forward neural network. Experiments reveal two main findings: (i) combining molecular structure information from SMILES with drug mono side-effect data substantially improves DDI prediction performance; and (ii) relying solely on chemical embeddings is insufficient.

## A Technical Appendices and Supplementary Material

Technical appendices with additional results, figures, graphs and proofs may be submitted with the paper submission before the full submission deadline (see above), or as a separate PDF in the ZIP file below before the supplementary material deadline. There is no page limit for the technical appendices.

### NeurIPS Paper Checklist

The checklist is designed to encourage best practices for responsible machine learning research, addressing issues of reproducibility, transparency, research ethics, and societal impact. Do not remove the checklist: **The papers not including the checklist will be desk rejected**. The checklist should follow the references and follow the (optional) supplemental material. The checklist does NOT count towards the page limit.

Please read the checklist guidelines carefully for information on how to answer these questions. For each question in the checklist:

- You should answer [Yes] , [No] , or [NA] .
- [NA] means either that the question is Not Applicable for that particular paper or the relevant information is Not Available.
- Please provide a short (1–2 sentence) justification right after your answer (even for NA).

#### The checklist answers are an integral part of your paper submission

They are visible to the reviewers, area chairs, senior area chairs, and ethics reviewers. You will be asked to also include it (after eventual revisions) with the final version of your paper, and its final version will be published with the paper.

The reviewers of your paper will be asked to use the checklist as one of the factors in their evaluation. While “[Yes] “ is generally preferable to “[No] “, it is perfectly acceptable to answer “[No] “ provided a proper justification is given (e.g., “error bars are not reported because it would be too computationally expensive” or “we were unable to find the license for the dataset we used”). In general, answering “[No] “ or “[NA] “ is not grounds for rejection. While the questions are phrased in a binary way, we acknowledge that the true answer is often more nuanced, so please just use your best judgment and write a justification to elaborate. All supporting evidence can appear either in the main paper or the supplemental material, provided in appendix. If you answer [Yes] to a question, in the justification please point to the section(s) where related material for the question can be found.

IMPORTANT, please:

- **Delete this instruction block, but keep the section heading “NeurIPS Paper Checklist”**,
- **Keep the checklist subsection headings, questions/answers and guidelines below**.
- **Do not modify the questions and only use the provided macros for your answers**.

##### 1. Claims

Question: Do the main claims made in the abstract and introduction accurately reflect the paper’s contributions and scope?

Answer: **[TODO]**

Justification: **[TODO]**

Guidelines:

- The answer NA means that the abstract and introduction do not include the claims made in the paper.
- The abstract and/or introduction should clearly state the claims made, including the contributions made in the paper and important assumptions and limitations. A No or NA answer to this question will not be perceived well by the reviewers.
- The claims made should match theoretical and experimental results, and reflect how much the results can be expected to generalize to other settings.
- It is fine to include aspirational goals as motivation as long as it is clear that these goals are not attained by the paper.

##### 2. Limitations

Question: Does the paper discuss the limitations of the work performed by the authors?

Answer: **[TODO]**

Justification: **[TODO]**

Guidelines:

- The answer NA means that the paper has no limitation while the answer No means that the paper has limitations, but those are not discussed in the paper.
- The authors are encouraged to create a separate “Limitations” section in their paper.
- The paper should point out any strong assumptions and how robust the results are to violations of these assumptions (e.g., independence assumptions, noiseless settings, model well-specification, asymptotic approximations only holding locally). The authors should reflect on how these assumptions might be violated in practice and what the implications would be.
- The authors should reflect on the scope of the claims made, e.g., if the approach was only tested on a few datasets or with a few runs. In general, empirical results often depend on implicit assumptions, which should be articulated.
- The authors should reflect on the factors that influence the performance of the approach. For example, a facial recognition algorithm may perform poorly when image resolution is low or images are taken in low lighting. Or a speech-to-text system might not be used reliably to provide closed captions for online lectures because it fails to handle technical jargon.
- The authors should discuss the computational efficiency of the proposed algorithms and how they scale with dataset size.
- If applicable, the authors should discuss possible limitations of their approach to address problems of privacy and fairness.
- While the authors might fear that complete honesty about limitations might be used by reviewers as grounds for rejection, a worse outcome might be that reviewers discover limitations that aren’t acknowledged in the paper. The authors should use their best judgment and recognize that individual actions in favor of transparency play an important role in developing norms that preserve the integrity of the community. Reviewers will be specifically instructed to not penalize honesty concerning limitations.

##### 3. Theory assumptions and proofs

Question: For each theoretical result, does the paper provide the full set of assumptions and a complete (and correct) proof?

Answer: **[TODO]**

Justification: **[TODO]**

Guidelines:

- The answer NA means that the paper does not include theoretical results.
- All the theorems, formulas, and proofs in the paper should be numbered and cross-referenced.
- All assumptions should be clearly stated or referenced in the statement of any theorems.
- The proofs can either appear in the main paper or the supplemental material, but if they appear in the supplemental material, the authors are encouraged to provide a short proof sketch to provide intuition.
- Inversely, any informal proof provided in the core of the paper should be complemented by formal proofs provided in appendix or supplemental material.
- Theorems and Lemmas that the proof relies upon should be properly referenced.

##### 4. Experimental result reproducibility

Question: Does the paper fully disclose all the information needed to reproduce the main experimental results of the paper to the extent that it affects the main claims and/or conclusions of the paper (regardless of whether the code and data are provided or not)?

Answer: **[TODO]**

Justification: **[TODO]**

Guidelines:

- The answer NA means that the paper does not include experiments.
- If the paper includes experiments, a No answer to this question will not be perceived well by the reviewers: Making the paper reproducible is important, regardless of whether the code and data are provided or not.
- If the contribution is a dataset and/or model, the authors should describe the steps taken to make their results reproducible or verifiable.
- Depending on the contribution, reproducibility can be accomplished in various ways. For example, if the contribution is a novel architecture, describing the architecture fully might suffice, or if the contribution is a specific model and empirical evaluation, it may be necessary to either make it possible for others to replicate the model with the same dataset, or provide access to the model. In general. releasing code and data is often one good way to accomplish this, but reproducibility can also be provided via detailed instructions for how to replicate the results, access to a hosted model (e.g., in the case of a large language model), releasing of a model checkpoint, or other means that are appropriate to the research performed.
- While NeurIPS does not require releasing code, the conference does require all submissions to provide some reasonable avenue for reproducibility, which may depend on the nature of the contribution. For example
  a. If the contribution is primarily a new algorithm, the paper should make it clear how to reproduce that algorithm.
  b. If the contribution is primarily a new model architecture, the paper should describe the architecture clearly and fully.
  c. If the contribution is a new model (e.g., a large language model), then there should either be a way to access this model for reproducing the results or a way to reproduce the model (e.g., with an open-source dataset or instructions for how to construct the dataset).
  d. We recognize that reproducibility may be tricky in some cases, in which case authors are welcome to describe the particular way they provide for reproducibility. In the case of closed-source models, it may be that access to the model is limited in some way (e.g., to registered users), but it should be possible for other researchers to have some path to reproducing or verifying the results.

##### 5. Open access to data and code

Question: Does the paper provide open access to the data and code, with sufficient instructions to faithfully reproduce the main experimental results, as described in supplemental material?

Answer: **[TODO]**

Justification: **[TODO]**

Guidelines:

- The answer NA means that paper does not include experiments requiring code.
- Please see the NeurIPS code and data submission guidelines (https://nips.cc/public/guides/CodeSubmissionPolicy) for more details.
- While we encourage the release of code and data, we understand that this might not be possible, so “No” is an acceptable answer. Papers cannot be rejected simply for not including code, unless this is central to the contribution (e.g., for a new open-source benchmark).
- The instructions should contain the exact command and environment needed to run to reproduce the results. See the NeurIPS code and data submission guidelines (https://nips.cc/public/guides/CodeSubmissionPolicy) for more details.
- The authors should provide instructions on data access and preparation, including how to access the raw data, preprocessed data, intermediate data, and generated data, etc.
- The authors should provide scripts to reproduce all experimental results for the new proposed method and baselines. If only a subset of experiments are reproducible, they should state which ones are omitted from the script and why.
- At submission time, to preserve anonymity, the authors should release anonymized versions (if applicable).
- Providing as much information as possible in supplemental material (appended to the paper) is recommended, but including URLs to data and code is permitted.

##### 6. Experimental setting/details

Question: Does the paper specify all the training and test details (e.g., data splits, hyperparameters, how they were chosen, type of optimizer, etc.) necessary to understand the results?

Answer: **[TODO]**

Justification: **[TODO]**

Guidelines:

- The answer NA means that the paper does not include experiments.
- The experimental setting should be presented in the core of the paper to a level of detail that is necessary to appreciate the results and make sense of them.
- The full details can be provided either with the code, in appendix, or as supplemental material.

##### 7. Experiment statistical significance

Question: Does the paper report error bars suitably and correctly defined or other appropriate information about the statistical significance of the experiments?

Answer: **[TODO]**

Justification: **[TODO]**

Guidelines:

- The answer NA means that the paper does not include experiments.
- The authors should answer “Yes” if the results are accompanied by error bars, confidence intervals, or statistical significance tests, at least for the experiments that support the main claims of the paper.
- The factors of variability that the error bars are capturing should be clearly stated (for example, train/test split, initialization, random drawing of some parameter, or overall run with given experimental conditions).
- The method for calculating the error bars should be explained (closed form formula, call to a library function, bootstrap, etc.)
- The assumptions made should be given (e.g., Normally distributed errors).
- It should be clear whether the error bar is the standard deviation or the standard error of the mean.
- It is OK to report 1-sigma error bars, but one should state it. The authors should preferably report a 2-sigma error bar than state that they have a 96% CI, if the hypothesis of Normality of errors is not verified.
- For asymmetric distributions, the authors should be careful not to show in tables or figures symmetric error bars that would yield results that are out of range (e.g. negative error rates).
- If error bars are reported in tables or plots, The authors should explain in the text how they were calculated and reference the corresponding figures or tables in the text.

##### 8. Experiments compute resources

Question: For each experiment, does the paper provide sufficient information on the computer resources (type of compute workers, memory, time of execution) needed to reproduce the experiments?

Answer: **[TODO]**

Justification: **[TODO]**

Guidelines:

- The answer NA means that the paper does not include experiments.
- The paper should indicate the type of compute workers CPU or GPU, internal cluster, or cloud provider, including relevant memory and storage.
- The paper should provide the amount of compute required for each of the individual experimental runs as well as estimate the total compute.
- The paper should disclose whether the full research project required more compute than the experiments reported in the paper (e.g., preliminary or failed experiments that didn’t make it into the paper).

##### 9. Code of ethics

Question: Does the research conducted in the paper conform, in every respect, with the NeurIPS Code of Ethics https://neurips.cc/public/EthicsGuidelines?

Answer: **[TODO]**

Justification: **[TODO]**

Guidelines:

- The answer NA means that the authors have not reviewed the NeurIPS Code of Ethics.
- If the authors answer No, they should explain the special circumstances that require a deviation from the Code of Ethics.
- The authors should make sure to preserve anonymity (e.g., if there is a special consideration due to laws or regulations in their jurisdiction).

##### 10. Broader impacts

Question: Does the paper discuss both potential positive societal impacts and negative societal impacts of the work performed?

Answer: **[TODO]**

Justification: **[TODO]**

Guidelines:

- The answer NA means that there is no societal impact of the work performed.
- If the authors answer NA or No, they should explain why their work has no societal impact or why the paper does not address societal impact.
- Examples of negative societal impacts include potential malicious or unintended uses (e.g., disinformation, generating fake profiles, surveillance), fairness considerations (e.g., deployment of technologies that could make decisions that unfairly impact specific groups), privacy considerations, and security considerations.
- The conference expects that many papers will be foundational research and not tied to particular applications, let alone deployments. However, if there is a direct path to any negative applications, the authors should point it out. For example, it is legitimate to point out that an improvement in the quality of generative models could be used to generate deepfakes for disinformation. On the other hand, it is not needed to point out that a generic algorithm for optimizing neural networks could enable people to train models that generate Deepfakes faster.
- The authors should consider possible harms that could arise when the technology is being used as intended and functioning correctly, harms that could arise when the technology is being used as intended but gives incorrect results, and harms following from (intentional or unintentional) misuse of the technology.
- If there are negative societal impacts, the authors could also discuss possible mitigation strategies (e.g., gated release of models, providing defenses in addition to attacks, mechanisms for monitoring misuse, mechanisms to monitor how a system learns from feedback over time, improving the efficiency and accessibility of ML).

##### 11. Safeguards

Question: Does the paper describe safeguards that have been put in place for responsible release of data or models that have a high risk for misuse (e.g., pretrained language models, image generators, or scraped datasets)?

Answer: **[TODO]**

Justification: **[TODO]**

Guidelines:

- The answer NA means that the paper poses no such risks.
- Released models that have a high risk for misuse or dual-use should be released with necessary safeguards to allow for controlled use of the model, for example by requiring that users adhere to usage guidelines or restrictions to access the model or implementing safety filters.
- Datasets that have been scraped from the Internet could pose safety risks. The authors should describe how they avoided releasing unsafe images.
- We recognize that providing effective safeguards is challenging, and many papers do not require this, but we encourage authors to take this into account and make a best faith effort.

##### 12. Licenses for existing assets

Question: Are the creators or original owners of assets (e.g., code, data, models), used in the paper, properly credited and are the license and terms of use explicitly mentioned and properly respected?

Answer: **[TODO]**

Justification: **[TODO]**

Guidelines:

- The answer NA means that the paper does not use existing assets.
- The authors should cite the original paper that produced the code package or dataset.
- The authors should state which version of the asset is used and, if possible, include a URL.
- The name of the license (e.g., CC-BY 4.0) should be included for each asset.
- For scraped data from a particular source (e.g., website), the copyright and terms of service of that source should be provided.
- If assets are released, the license, copyright information, and terms of use in the package should be provided. For popular datasets, paperswithcode.com/datasets has curated licenses for some datasets. Their licensing guide can help determine the license of a dataset.
- For existing datasets that are re-packaged, both the original license and the license of the derived asset (if it has changed) should be provided.
- If this information is not available online, the authors are encouraged to reach out to the asset’s creators.

##### 13. New assets

Question: Are new assets introduced in the paper well documented and is the documentation provided alongside the assets?

Answer: **[TODO]**

Justification: **[TODO]**

Guidelines:

- The answer NA means that the paper does not release new assets.
- Researchers should communicate the details of the dataset/code/model as part of their submissions via structured templates. This includes details about training, license, limitations, etc.
- The paper should discuss whether and how consent was obtained from people whose asset is used.
- At submission time, remember to anonymize your assets (if applicable). You can either create an anonymized URL or include an anonymized zip file.

##### 14. Crowdsourcing and research with human subjects

Question: For crowdsourcing experiments and research with human subjects, does the paper include the full text of instructions given to participants and screenshots, if applicable, as well as details about compensation (if any)?

Answer: **[TODO]**

Justification: **[TODO]**

Guidelines:

- The answer NA means that the paper does not involve crowdsourcing nor research with human subjects.
- Including this information in the supplemental material is fine, but if the main contribution of the paper involves human subjects, then as much detail as possible should be included in the main paper.
- According to the NeurIPS Code of Ethics, workers involved in data collection, curation, or other labor should be paid at least the minimum wage in the country of the data collector.

##### 15. Institutional review board (IRB) approvals or equivalent for research with human subjects

Question: Does the paper describe potential risks incurred by study participants, whether such risks were disclosed to the subjects, and whether Institutional Review Board (IRB) approvals (or an equivalent approval/review based on the requirements of your country or institution) were obtained?

Answer: **[TODO]**

Justification: **[TODO]**

Guidelines:

- The answer NA means that the paper does not involve crowdsourcing nor research with human subjects.
- Depending on the country in which research is conducted, IRB approval (or equivalent) may be required for any human subjects research. If you obtained IRB approval, you should clearly state this in the paper.
- We recognize that the procedures for this may vary significantly between institutions and locations, and we expect authors to adhere to the NeurIPS Code of Ethics and the guidelines for their institution.
- For initial submissions, do not include any information that would break anonymity (if applicable), such as the institution conducting the review.

##### 16. Declaration of LLM usage

Question: Does the paper describe the usage of LLMs if it is an important, original, or non-standard component of the core methods in this research? Note that if the LLM is used only for writing, editing, or formatting purposes and does not impact the core methodology, scientific rigorousness, or originality of the research, declaration is not required.

Answer: **[TODO]**

Justification: **[TODO]**

Guidelines:

- The answer NA means that the core method development in this research does not involve LLMs as any important, original, or non-standard components.
- Please refer to our LLM policy (https://neurips.cc/Conferences/2025/LLM) for what should or should not be described.

